# Structural mutations set an equilibrium non-coding genome fraction

**DOI:** 10.1101/2025.02.03.636187

**Authors:** Juliette Luiselli, Paul Banse, Olivier Mazet, Nicolas Lartillot, Guillaume Beslon

## Abstract

Non-coding genome size evolution is poorly understood. While some fraction of non-coding DNA has arguably a regulatory function, a large part does not seem to have a detectable impact on any phenotypic trait. The abundance of non-functional DNA in genomes, observed across the Tree of Life, challenges a purely adaptationist explanation. Several non-adaptive theories have been proposed to explain its presence and identify its determinants, emphasizing either the mutational processes or the mutational hazard entailed by non-coding and non-functional DNA. However, those theories have not yet been integrated into a single framework, and the exact nature of the mutational hazard is not yet fully understood. In this work, we propose a simple mathematical model of genome size evolution. The model shows how the non-coding fraction of the genome is shaped by two factors: unavoidable biases in the neutrality of the different mutation types (adding base pairs is more likely to be neutral than removing some), and the robustness selection imposed by the mere existence of structural mutations (larger genomes are more prone to double-strand breaks that can initiate structural mutations, imposing a second-order selection on robustness). Together, these two factors ensure the existence of an equilibrium non-coding fraction. We show that this equilibrium depends solely on mutation biases and the product of population size and mutation rate.

## Introduction

Genome size varies greatly throughout the Tree of Life: from 10^5^ base pairs (bp) for some bacteria [37], to more than 10^11^ bp for some plants [34]. Coding sequences contribute to this variation through adaptive changes, but some parts of the genome seem devoid of phenotypic function and yet are highly variable in size [21]. While non-coding DNA contains functional sequences, including regulatory regions [38], large stretches seem to bear no function whatsoever. This “junk” DNA [32, 8, 33, 10] is ubiquitous in all domains of life, regardless of genome sizes [1, 14]. However, there is currently no consensus on the reasons behind the existence and maintenance of junk DNA [10].

In this work, we address the determinants of the amount of non-coding non-functional DNA. Several hypotheses have been proposed to address these issues, notably reviewed in [6].

In adaptive hypotheses, genome size itself is under selection due to its phenotypic impact on *e*.*g*. nucleus size or replication time [28]. In this view, genome size would be selectively limited [18, 3]. Furthermore, the position of genes relative to each other or the centromere influences their expression [9]. As such, it represents a potential selective pressure on the amount of intergenic DNA [13]. However, it can be argued that the variation in the proportion of non-coding DNA between species might be too high to be explained by these mechanisms [35, 6]. More fundamentally, there is little direct evidence that selection induced by these phenotypes is strong enough to modulate the fate of mutations changing genome size.

Non-adaptive hypotheses have also been developed to decipher the mechanisms by which non-coding DNA could vary and stabilize. First, mutational explanations emphasize the impact of mutational patterns on the longterm evolution of genome size. In particular, the mutational equilibrium hypothesis (MEH)[35] suggests that two different mutational biases of opposite directions —a negative bias on short indels and a positive long insertion/deletion bias that decreases with genome size —could mechanistically explain the existence of an equilibrium genome size. The equilibrium itself would be modulated between species by the variation in the strength of those biases.

The mutational hazard hypothesis (MHH)[27], on the other hand, proposes an explanation in terms of *fixation* biases acting on mutations, related to second-order selective effects. According to the MHH, the non-coding genome expands by mutation, drift, and the insertion of selfish elements. However, this expansion increases the number of targets for deleterious mutations —*e*.*g*., such as gain-of-function mutations or loss of accurate splicing (ref Lynch book). In other words, non-coding DNA presents a mutational liability. As a result, genome expansion entails a slight selective cost, which could provide a sufficient force counteracting the growth of genome size [27, 24]. The efficacy of this selective force is inversely related to effective population size, while the intensity of the force itself is directly proportional to the mutation rate. Thus, genome size should be inversely correlated with each of these two factors [24, 20].

Both theories receive support from some observations [46, 19, 42, 31, 7, 41, 23], but are also challenged by others [2, 40, 30, 29]. Importantly, they are not mutually exclusive, as a combination of mutational biases and second-order selective effects due to the mutational liability of non-functional DNA could act together to determine an equilibrium genome size. This calls for an integrated explanation for what determines the amount and variation of non-coding DNA in genomes. In this direction, previous studies on simulated data [4, 23] suggest that structural mutations, *i*.*e*. chromosomal rearrangements or more generally any mutation larger than 50 bp, could be a key element linking both the MEH and the MHH. Indeed, structural mutations significantly affect genome size and are also a huge mutational liability in themselves due to their large-scale effect. It is therefore essential to examine their impact on genome size evolution.

Here, we propose a minimal probabilistic model of genome evolution, with the following assumptions: (1) genomes are composed of a coding component made of essential genes and a non-coding component that has strictly no phenotypic effect; (2) mutations occur at random uniformly over the genome. Our analysis of this model reveals non-trivial patterns: (1) structural mutations do not have the same probability of being neutral and this results in a trend towards increasing genome size; (2) as larger genomes are more susceptible to double-strand breaks —and thus to structural mutations —, changes in genome size change the probability of future, possibly lethal, structural mutations; (3) this increased risk of having a larger genome modulates the fixation probability of structural mutations in a way that favors deletions over insertions or duplications. Together, these mechanisms ensure a stable evolutionary equilibrium for non-coding genome size. More precisely, the equilibrium non-coding *fraction* depends on the product of the effective population size and mutation rate of a species (*N*_*e*_ *× µ*), while the non-coding *size* is determined by this product plus the coding architecture of the genome (coding size and distribution). Notably, the equilibrium is a robust outcome of our model, even in the presence of mutational biases towards insertions or deletions: arbitrary mutational biases merely shift the equilibrium.

Altogether, our model integrates key aspects of the MEH and MHH to provide a general mechanistic explanation for genome size evolution. It highlights structural mutations as a major mutational hazard susceptible to driving genome size evolution under general conditions.

## Model and Results

### Model overview and existence of an equilibrium non-coding genome size

To address the question of non-coding genome size evolution, we study the effect of mutations on a population of *N* individuals with simplified, circular genomes.

As shown in Fig. 1, we consider a circular haploid genome of length *L* base pairs (bp), composed of *g* coding segments (and thus *g* non-coding segments). Let us note the number of non-coding base pairs *z*_nc_ and the number of coding base pairs *z*_*c*_. We have *z*_c_ + *z*_nc_ = *L*. We assume that:

**Figure 1.**
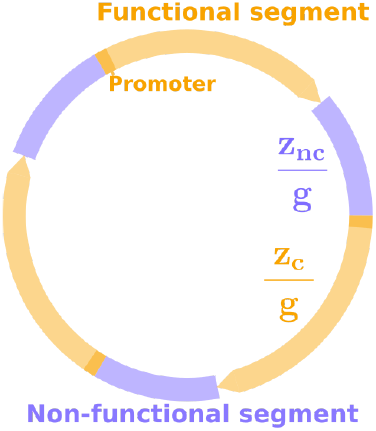
Representation of a genome, with *g* = 3. Non-coding segments are of the same size *z*_nc_ */g*, and coding segments are of the same size *z*_c_*/g*. Each coding segment starts with a promoter that can create a new coding segment if duplicated.

- Coding segments are of the size 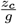 and represent “genes”. Genes are non-overlapping and all oriented in the same direction. The non-coding segments are equally distributed between the *g* genes and are each of size 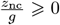. We assume this remains true after any change in non-coding size, as neutral inversions will reshuffle the genome. This ensures that a genome can be fully described with just *g, z*_c_ and *z*_nc_.
- Deleting any base of a gene inactivates it and is always lethal. Genes are assumed to have a promoter, here represented by their first base. As a result, a partial duplication not including this first base is not expressed and is thus neutral, *i*.*e*. it does not affect viability. Conversely, a partial or complete duplication including the promoter results in a new expressed gene and is also lethal.
- Non-lethal mutations are assumed to be perfectly neutral for the viability of the individual. Thus, fitness is binary: it is either 1 or 0.

Different types of mutations occur at different mutation rates. We note *µ* the basal per base mutation rate of the organism, and *λ*_*i*_*µ* is the per base mutation rate for mutation type *i*. Throughout the manuscript, we analyze the evolution of the non-coding genome size *z*_nc_, under the assumption that the coding genome size *z*_c_ and the number of coding segments *g* remain fixed.

### Neutral genome growth

We compute the probability of different types of mutations to be neutral and fixed in a population of size *N* . In the following, a mutation is said to be neutral when it does not affect the coding genome and thus does not alter the viability of the individual.

For the sake of clarity, we consider here (sections and) a simple version of the model including only two types of structural mutations: duplications (dupl) and deletions (del), occurring at the same per bp rate (*λ*_dupl_ = *λ*_del_ = 1). A duplication copies a random segment of the genome and inserts it elsewhere, while a deletion removes a random segment of the genome. The breakpoints are chosen uniformly at random on the genome, such that both mutations have the same size distribution and the expected change of size of the genome upon one mutation is 0.

We calculate the probability *ν* for each of these two mutations to be neutral (in terms of viability), recalling that duplicating a promoter, inserting a segment within a gene, or deleting any base of a gene is always deleterious. Detailed computations are provided in Supplementary Materials (section S1).

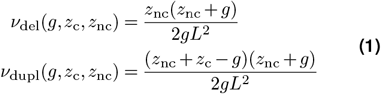

Notably, we have : 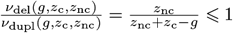,as *z*_c_ is obviously much larger than *g*. Thus, duplications are more often neutral than deletions. Similarly, we show that neutral duplications are also on average larger than neutral deletions (see Supplementary Materials S6). As illustrated by Fig. 2A, we can also consider the probability for a mutation of a given size *k* to be neutral. It highlights that increasing *z*_nc_ increases both the probability for mutations of a given size to be neutral and the range of possible neutral mutations (note that, above a certain size, mutations are always lethal due to constraints from the genome architecture: larger mutations would necessarily delete part of a gene or duplicate a promoter). Consequently, genomes should grow indefinitely if we assume that only neutral mutations are fixed with an equal probability. However, a mutation that is neutral in terms of fitness for the individual is not necessarily neutral in terms of fitness for the lineage. The next subsection will explore the effect of this second-order selective force.

**Figure 2.**
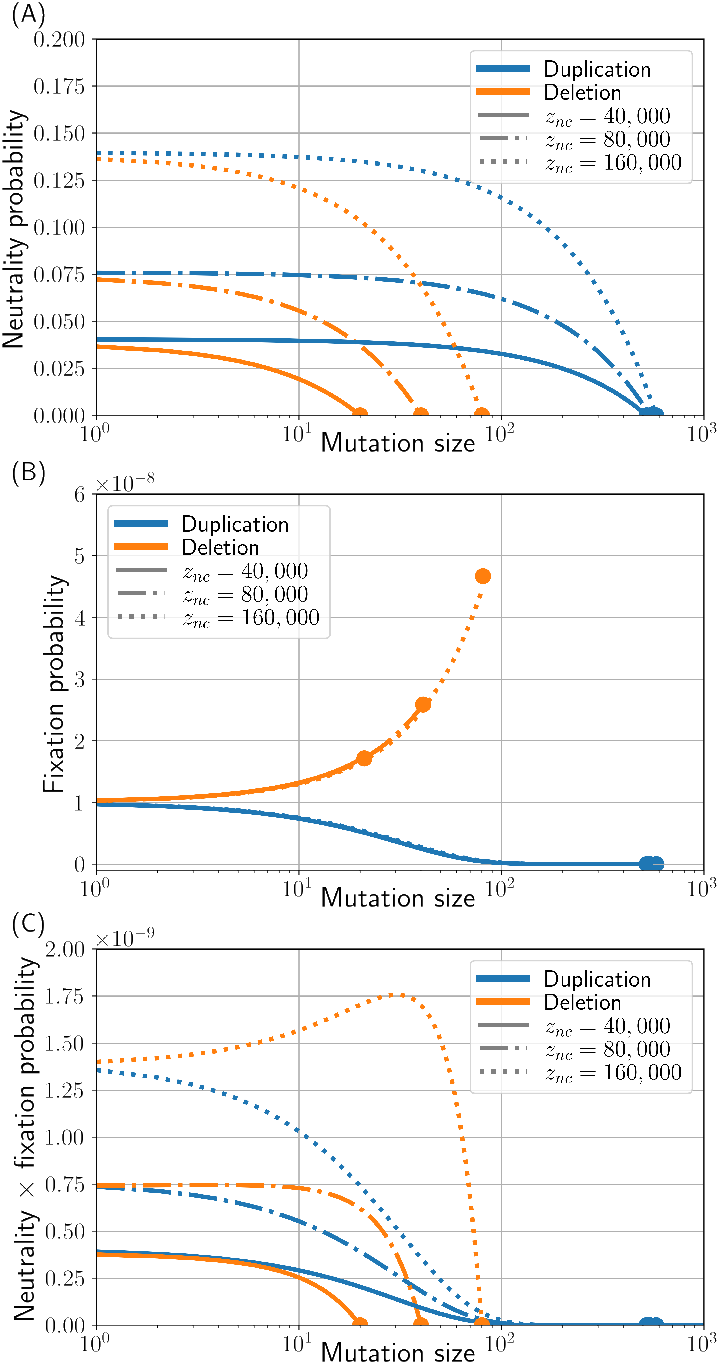
Effect of mutation type and genome size on neutrality and fixation. The dots mark points above which mutations cannot be neutral nor fixed due to constraints from the genome architecture. **(A)** Probability of a duplication (blue) or a deletion (orange) to not affect the coding genome for different mutation sizes and non-coding sizes. The number of genes *g* is fixed at 2, 000, and the coding genome size at *z*_c_ = 1, 000, 000 bp. Mutations are more likely to be neutral in bigger and more non-coding genomes, and neutral duplications are larger and more frequent than neutral deletions. **(B)** Probability of fixation of a neutral duplication (blue) or a neutral deletion (orange) for different mutation sizes and non-coding sizes. The number of genes *g* is fixed at 2, 000, the coding genome size at *z*_c_ = 1, 000, 000 bp, and the population size is *N* = 10^8^. **(C)** Probability of being neutral *and* fixed for different mutation sizes and different non-coding sizes. The number of genes *g* is fixed at 2, 000, the coding genome size at *z*_c_ = 1, 000, 000 bp, and the population size is *N* = 10^8^. For the shortest non-coding size (plain line), duplications are more often neutral and fixed than deletions for any mutation size, indicating that the non-coding genome size would increase. On the contrary, for the biggest non-coding size (dotted line) deletions are more often neutral and fixed than duplications for any mutation size, indicating that the non-coding genome size would decrease: there must be an equilibrium non-coding size between these values.

### Robustness selection

By definition, a neutral duplication or deletion does not change the viability of an individual. However, it changes the non-coding genome size *z*_nc_. Now, the probability for a mutation to be neutral depends on *z*_nc_ (see Eq. 1), and so a neutral mutation changes the probability for future mutations to also be neutral. Changing the genome size also changes the probability for a mutation to occur at replication, as bigger genomes will naturally undergo more mutations for the same per base mutation rate. Therefore, mutations that are neutral in terms of their immediate effect on the viability still change the probability for the individual to have future offspring that are equally fit —their robustness. For the rest of the manuscript, we call the *effective fitness f*_*e*_ of an individual the average fitness of its potential offspring. This can also be viewed as the fecundity of an individual once the viability of the offspring has been taken into account. In our model, supposing that at most one mutation of each type can occur upon replication, we have:

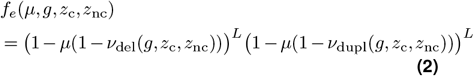

Since genome size modifies the effective fitness and affects a lineage survival probability in the long term, it can be selected. In particular, while increasing the non-coding genome size *z*_nc_ increases the probability for mutations to be neutral (see Fig. 2A), it also increases the probability for a mutation to happen. As a result, the effective fitness *f*_*e*_ actually decreases as the non-coding genome size increases (see Fig. 3), and selection can then act against the genome size increase described in paragraph A1.

**Figure 3.**
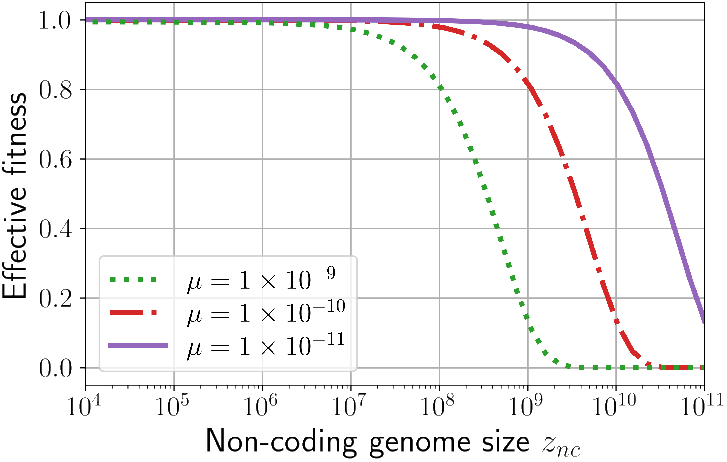
Effective fitness *f*_*e*_ for different non-coding sizes *z*_nc_ and different mutation rates *µ*. Genome architecture is fixed at *z*_c_ = 1, 000, 000 and *g* = 2, 000, and *λ*_del_ = *λ*_dupl_ = 1. Notably, the effective fitness decreases with both *z*_nc_ and *µ*.

We characterize this effect more precisely using a population genetics argument. We consider a haploid population of wild-type individuals of size *N* in which a mutant appears and bears a neutral mutation that adds *k* bases to its non-coding genome, with *k* ∈ ℤ^∗^. *k* can be either positive (duplication) or negative (deletion). We consider that the population follows a Wright-Fisher model [12, 45], and we compute the probability for this mutant to go to fixation [39]:

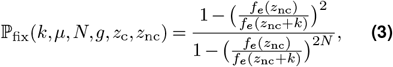

where we note *f*_*e*_(*µ, g, z*_c_, *z*_nc_) simply *f*_*e*_(*z*_nc_), as we consider that other parameters are fixed. As illustrated by Fig. 2B, mutations that increase genome size are less likely to be fixed than mutations that decrease genome size. This is the direct consequence of an increase in genome size being tied to an increase in the per genome mutation rate, and hence a decrease in effective fitness (see Fig. 3). Hence, while neutral duplications are more frequent and larger than neutral deletions, they are also more rarely fixed. When considering the combination of these two tendencies (the opposing biases in the immediate probability of being lethal and in the ultimate probability of being fixed), we can see that the shortest genomes are more likely to fix neutral duplications while the longest genomes are more likely to fix neutral deletions, as demonstrated in Fig. 2C. This intuitively results in an equilibrium genome size at which the two effects cancel out.

### Computing the equilibrium non-coding genome size

To formalize this equilibrium genome size, we compute the average contribution of duplications (*δ*_dupl_) and deletions (*δ*_del_) to changes in non-coding genome size in the population. *δ*_dupl_ and *δ*_del_ are expressed in bp per generation per mutation event and represent the average length of fixed mutations per time unit. They are computed under the origination-fixation approximation, meaning there is no clonal interference, and we consider the probability for each mutation individually to go to fixation in the absence of any other mutant in the population. Each *δ* thus depends on the mutation’s probability of being neutral, its size, and its fixation probability:

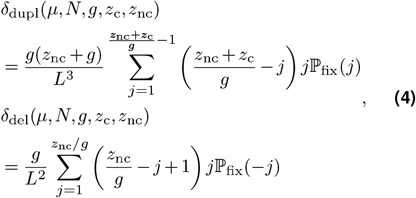

where we denote ℙ_fix_(*k, µ, N, g, z*_c_, *z*_nc_) as ℙ_fix_(*k*), as other parameters are supposed fixed. Detailed derivations are presented in the Supplementary Materials, section S2. From Eq. 4, we can derive the bias towards increasing or decreasing genome size as the ratio between the sum of the contributions of all deletions over the sum of the contributions of all duplications for a given genome size, population size, and mutation rate.

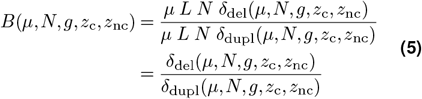

The non-coding genome size is at equilibrium when *B* = 1 (see Fig. 4). When the bias is above 1, deletions contribute more to genome size changes and the non-coding proportion shrinks. On the other hand, when the bias is below 1, duplications contribute more to genome size change and the non-coding proportion increases.

**Figure 4.**
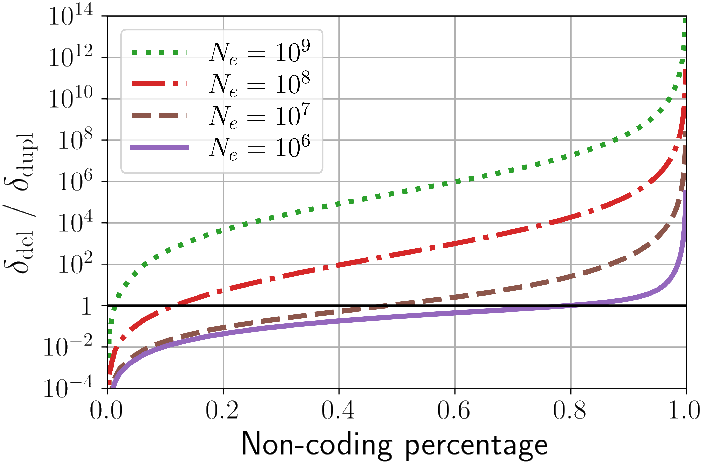
Measured bias for different non-coding proportions. Genome architecture is fixed at *z*_c_ = 1, 000, 000 and *g* = 2, 000, the mutation rate is fixed at *µ* = 1 *×* 10^−10^ and *λ*_del_ = *λ*_dupl_ = 1. *z*_nc_ varies in a logspace from 10^3^ to 10^9^, and four different values of *N* are depicted, showing a progression in the equilibrium non-coding percentage. The black horizontal line shows the equilibrium at *B* = 1.

### Joint impact of population size and mutation rate

*B* is a function of the genome architecture (*z*_c_, *z*_nc_, and *g*), the population size *N*, the mutation rate *µ*. However, we can show that *B* depends on *N* and *µ* only through their product, as previously observed in simulation data [23].

Indeed, *N* and *µ* only appear in *f*_*e*_ (Eq. 2) and ℙ_*fix*_ (Eq. 3). Let us start with the expression of the effective fitness *f*_*e*_, and consider that the mutation rate *µ* is negligible compared to 1 (*µ* ≪ 1).

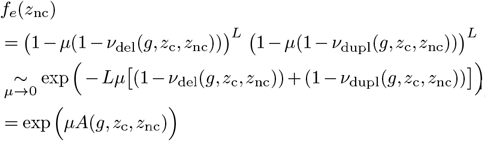

Where *A*(*g, z*_c_, *z*_nc_) = −(*z*_c_ + *z*_nc_)(1 − *ν*_del_(*g, z*_c_, *z*_nc_)) +(1 − *ν* (*g, z*, *z*)) *<* 0. Then, the ratio of effective fitnesses used in the computation of ℙ_*fix*_ (Eq. 3) can be written as:

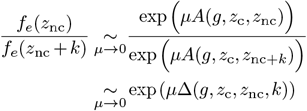

Where Δ(*g, z*_c_, *z*_nc_, *k*) = *A*(*g, z*_c_, *z*_nc_) − *A*(*g, z*_c_, *z*_nc_ + *k*) is a function that depends solely on genome architecture (*z*_c_, *z*_nc_ and *g*) and mutation size *k*. The probability of fixation of a mutation changing the genome size by *k* (positive or negative) is thus:

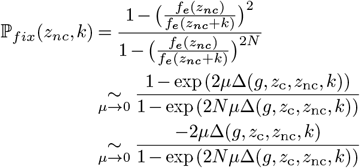

ℙ_*fix*_ appears to be a function of *µ, N × µ* and other parameters. Thus, both *δ*_dupl_ and *δ*_del_ can also be written as *µ* times a function of *N × µ*. Since 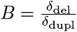,the *µ*s cancel out and *N* and *µ* always appear in the form of a product in *B*. Given a fixed coding size *z*_c_ and a number of coding segments *g* (*i*.*e*. a fixed coding architecture), *N* and *µ* have therefore a similar impact on the equilibrium non-coding size. This can be illustrated by a numerical exploration of the relative effects of *N* and *µ* (Supplementary Materials S3).

An alternative way to see this result is by noting that *µ*Δ(*g, z*_c_, *z*_nc_, *k*) is the selection coefficient associated with the effective fitness: 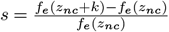.Indeed, in the limit *µ* → 0 and 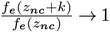,and thus *s* → 0, we have *s* ∼ ln 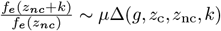.The mutation-selection-drift equilibrium, as a general rule, depends only on relative, not absolute, mutation rates (thus here, on the mutational bias). In addition, it depends on the various selective effects implicated in it only through their scaled selection coefficients. Here, *Ns*∼ *Nµ*Δ(*g, z*_c_, *z*_nc_, *k*), and thus, in the end, the mutation-selection-drift equilibrium depends on *N* and *µ* only through their product.

All these results show that an equilibrium non-coding genome size exists and depends on the coding genome architecture (*g* and *z*_c_) and on the product *N × µ*.

### Necessary condition for the existence of the equilibrium

So far, we only considered one type of mutation: structural variations. Other types of mutation change genome size and one could ask whether they would lead to a similar equilibrium. In particular, short indels (*<* 50 bp) can also add or remove bases to the genome and contribute to genome size changes, although less abruptly than structural variants. Most interestingly, if we replicate our model with only short indels (see Supplementary Materials sections S1 and S2), we don’t observe an equilibrium genome size, except under very specific conditions (for extremely large population size and starting from a small enough genome). In all other cases, *δ*_indel_+ *> δ*_indel_− and so, in the absence of a sufficiently strong mutational bias in favor of deletions, indels induce an infinite growth of the non-coding size (section 4), confirming previous observations [4]. Indeed, except in very specific ranges of parameters (see Supplementary Material section 4), indels do not create a selection for shorter genomes on their own: although they are more numerous as genome size increases, they are also more often neutral due to their size being limited, and so their effect is more likely to be limited to non-coding parts of the genome. On the opposite, structural variations, being driven by double-strand breaks, conserve their mutational liability when non-coding genome size increases. This makes them a necessary component to observe a pervasive genome size equilibrium.

### Expanded model of non-coding genome size evolution

Although the existence of an equilibrium specifically requires the presence of structural mutations, other types of mutations, with possibly different mutation rates, could contribute to genome size —directly (by changing the amount of non-coding sequences) or indirectly (due to their intrinsic mutational liability). To account for this, we added four types of mutations to our mathematical model: point mutations (pm), inversions (inv), small insertions (indel^+^), small deletions (indel^−^). Each mutation type *i* has its own mutation rate *λ*_*i*_*µ*. The probability of being neutral for all these mutations, and the average contribution to changes in genome size for indels, is presented in the Supplementary Materials (sections S1 and S2).

Naturally, these mutations and biases displace the equilibrium value of our model, as they change both the robustness of the genomes and the probability of removing or adding new bases. However, Fig. 5 shows that genome size (or equivalently the non-coding fraction) is always a decreasing function of *Nµ*, whatever the underlying mutational bias. Notably, when there is a deletion bias, the non-coding fraction remains bounded for all values of *Nµ*. The upper bound is the asymptotic value reached when *Nµ*→ 0, and it is smaller for more pronounced deletion biases. Thus, in this regime, not all coding fractions can be achieved by varying *Nµ*. On the other hand, when there is no bias or an insertion bias, the non-coding fraction diverges in the limit *Nµ* → 0, such that arbitrarily large non-coding fractions can be achieved with sufficiently small *Nµ*. However, there is always an equilibrium value for *Nµ >* 0.

**Figure 5.**
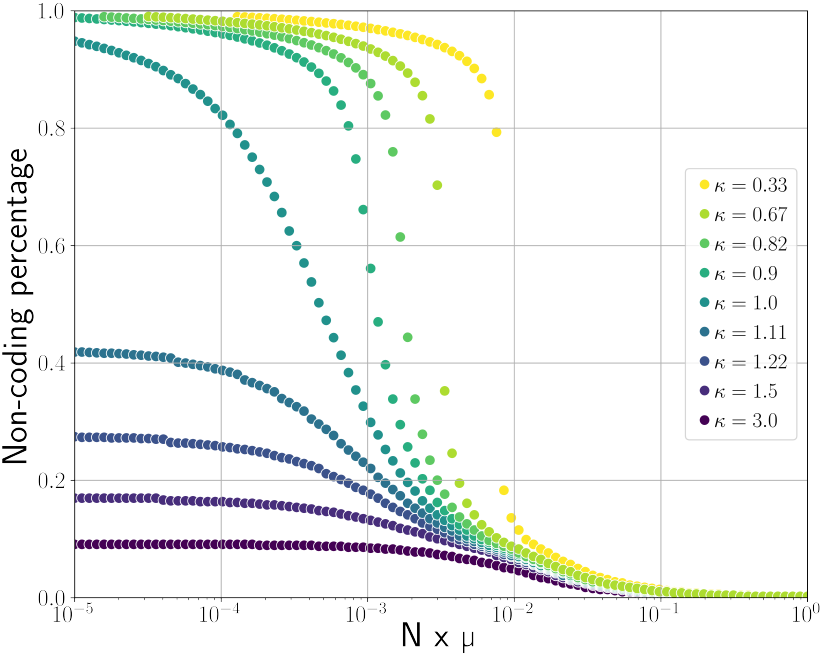
Predicted non-coding fractions for different values of *N × µ* using the expanded version of the model with six types of mutations. Two sets of equilibrium percentages were run: with *µ* = 10^−9^ and *N* varying from 10^4^ to 10^9^, and with *N* = 10^8^ and *µ* varying from 10^−13^ to 10^−8^. We note the deletion bias 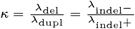. Note that we fix 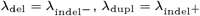, and these four *λi* sum to 4. Other parameters are fixed at *g* = 2, 000, *z*_c_ = 1, 000, 000, *l*_*m*_ = 50 and *λ*_pm_ = *λ*_inv_ = 1.

Notably, the addition of new mutations and the variations in the mutational bias do not suppress the existence of the equilibrium, as they do not fundamentally change the mechanisms at stake. The equilibrium is still determined by the product *N× µ* and the coding genome architecture (*z*_c_ and *g*), and the variations are always in the same direction: a higher population size or a higher mutation rate is associated with a lower non-coding fraction.

### Insights from biological data

Our model predicts that the *fraction* of the non-coding genome depends only on the compound parameter *N*_*e*_ *× µ*, with *µ* the structural mutation rate, as depicted by Fig. 5, as well as on the relative insertion versus deletion rates. The non-coding absolute size has more complicated dependencies, as it also depends on the coding architecture (see discussion). These predictions could in principle be tested against empirical data. However, spontaneous structural mutation rates are unknown, as most structural mutations are strongly deleterious, hence frequently purged by selection and notoriously difficult to observe and quantify [17]). Notwithstanding, a tentative comparison with empirical data is shown in Fig. 6, relying on nucleotide diversity to estimate *N*_*e*_*µ* and assuming a 1:1 ratio for structural versus point mutation rates (ratios of either 1:10 or 10:1 would shift empirical points to the left or the right, respectively, compared to the theoretical curves).

**Figure 6.**
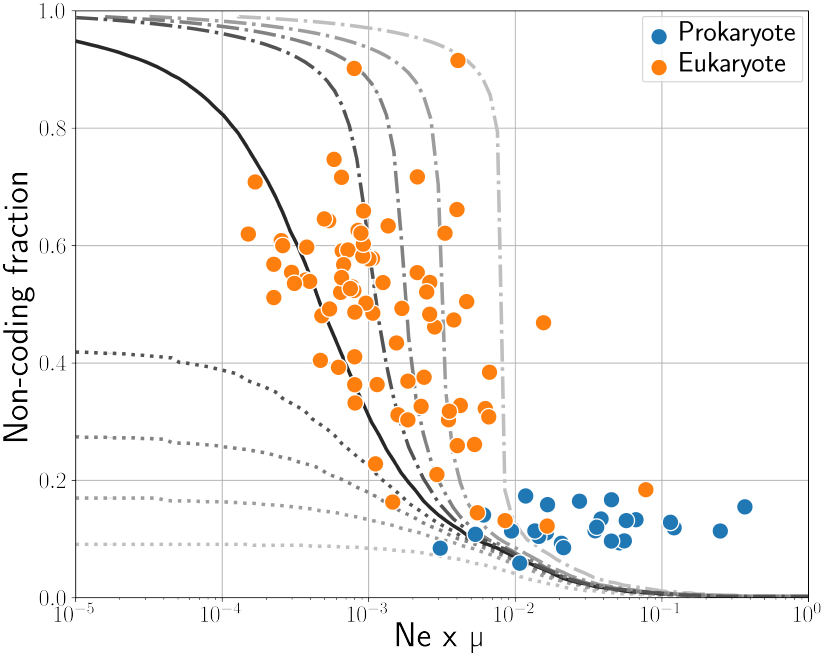
Non-coding fraction plotted against *N*_*e*_ *× µ* for 129 species from [26]. The mutation rate *µ* used here is the per generation per base substitution rate, which we assume to be correlated with the overall mutation rate. The gray lines show the equilibrium non-coding percentages predicted by our model for the same range of *N × µ* and different mutational biases *κ* (see Fig. 5).

With a structural mutation rate of this order of magnitude, our model globally predicts the overall trend of the distribution of non-coding fractions observed across species, as a function of nucleotide diversity. Thus, species with a high *N*_*e*_*× µ* present a lower coding fraction than species with a low *N*_*e*_ *× µ*. More precisely, eukaryotes are mostly located on the left of the figure and have both a higher non-coding fraction and a lower *N*_*e*_*× µ*, with a tendency to follow that relationship within them, while prokaryotes are on the right of the figure and present both a lower non-coding fraction and a higher *N*_*e*_*× µ*. Notably, the actual non-coding percentage of the prokaryotes shown here is higher than the one predicted by the model, but this is expected as our model assumes that the non-coding is purely non-functional, while the non-coding genome actually comprises regulatory RNAs and other functional sequences. This could indicate that most of the “non-coding” base pairs of prokaryotes have a phenotypic effect.

Altogether, and even if it is still far from a formal test, this comparison with empirical data gives an idea of the structural mutation rates for which second-order selection on genome rearrangements represents a key force preventing an infinite genome size growth. It also reproduces the global non-coding genome fraction variation patterns across cellular life. It shows that the potential role of structural mutations in non-coding genome size evolution should not be underestimated and deserves further investigation.

## Discussion

Our model reveals simple yet important evolutionary dynamics on genome size due to two opposite effects. On the one hand, duplications and insertions are more often neutral than deletions, implying a neutral bias towards genome size increase, a mechanism akin to the border selection effect proposed by [15, 22]. As an intuitive example, in the extreme case of a fully coding genome, it is still possible to neutrally insert a base between two genes, while no base can be removed. On the other hand, at constant phenotypical adaptation bigger genomes are counter-selected as they are more susceptible to structural mutations: lineages in which genomes get bigger are less likely to survive in the long term. This second-order selection for shorter genomes is imposed by the mere existence of structural mutations, which decrease the robustness of genomes when non-coding genome size increases —as previously conjectured [20]. With these two effects, and knowing only the genome’s current coding architecture (size and number of segments), the effective population size of the species, and the rates for the different mutation types, we can determine an equilibrium non-coding genome size towards which the species should be tending. Notably, this does not apply when structural mutations are absent from the model: with only indels, our model predicts that genomes are likely to grow indefinitely. Naturally, parameters not considered here could impact this equilibrium quantitatively. Most importantly, the presence of transposable elements, non-coding but functional DNA, horizontal transfers, and other mutational processes, such as recombination, would displace the equilibrium by affecting both the neutral mutational bias and the robustness of genomes. Yet, they would not suppress either of the two above-elaborated effects and so the existence of the equilibrium remains —as well as the direction of change in non-coding proportion caused by changes in *N* or *µ*. Similarly, we have assumed a uniform distribution for the sizes of mutations. Relaxing this assumption would also displace the equilibrium, although robustness selection would still operate, as long as the size of the structural mutations increases with genome size —which is intuitive as some species’ structural variants are longer than other species’ genomes [44]. Therefore, we expect the two effects we present here to be pervasive. In particular, while the hypotheses of our model are closer to a prokaryote-like genome (a single haploid circular chromosome), there is no reason for the general mechanism to not be true in the case of eukaryotes, and we can use it to compute the predicted non-coding percentage around eukaryote-like values of *N*_*e*_ and *µ* (as shown by Fig. 6).

As a first empirical confrontation of our theory, our comparison of the predictions of our model with empirical data (Fig. 6) results in a globally coherent and insightful picture for both eukaryotes and prokaryotes, as predictions could align with biological observations of non-coding fractions. In particular, our model predicts more variability of non-coding fractions in the eukaryotic parameter range, which lies around the steepest part of the curves (see Fig. 6). Conversely, prokaryotes, having a much larger *N*_*e*_ *× µ* product, are predicted to be much more stable around lower non-coding fractions. In that case, although a deletion bias in prokaryotes could exist, the *N*_*e*_ *× µ* values are in a range where the predicted non-coding percentage is only loosely affected by mutational biases. Mathematically, the high variation of non-coding percentages in eukaryotes could be explained by observing that the function *B* is flatter for lower values of *N*_*e*_ *× µ*, typical of eukaryotes (see Fig. 4). In these ranges, it is harder to reach the equilibrium non-coding value, as the bias towards losing or gaining bases at each generation is very low, hence allowing more variability of non-coding sizes at constant *N*_*e*_ × *µ* and constant mutational bias. Eukaryotes are supposed to be subjected to biases towards insertions [5], making it more complicated to compare data to our model. Finally, the results presented also depend on the relative rates of different types of mutations, which are largely unknown for structural mutations. Indeed, although they are frequently observed in all domains of life [36, 11], their spontaneous rate is very difficult to estimate due to their strong deleterious effect. As such, interpretations should be taken with caution. In short, our model proposes an explanation for the very different non-coding percentages observed in eukaryotes and prokaryotes, without postulating a difference in nature between these two types of organisms but only relying on the existence of structural mutations and the different values of *N*_*e*_ and *µ* for eukaryotes and prokaryotes.

Surprisingly, our results point out that non-coding genome *fractions* evolution is determined by the product *N*_*e*_*× µ* and the mutational bias, which, to our knowledge, is a new prediction. Although these factors have already been pointed out as the potential determinant of (non-coding) genome *size* [27, 46, 19, 35], we show that size also depends strongly on the coding architecture of a genome, with the latter probably highly driven by adaptation. This, as well as the fact that *N*_*e*_ and *µ* act jointly and should always be accounted for in the form of their product, could explain why some data are not aligned with the MHH [29, 2, 40, 30]. Consequently, further research should focus on non-coding fractions, or account for differences in coding architectures when comparing non-coding genome size and *N*_*e*_ and *µ* between species.

Finally, although we have assumed fixed mutation rates, in reality, mutation rates are themselves susceptible to evolve. According to the drift barrier model [43], mutation rates are under directional selection limited by random drift, such that they reach an evolutionary equilibrium that depends on *N*_*e*_. The selective force involved in this higher-order evolutionary process stems from the deleterious effects of new mutations on the offspring of the current generation [42, 43, 25]. Interestingly, in the case of structural mutations, this force is the same as the second-order selective force behind the mutational hazard (see Eq. 2). Thus, any mutational hazard also represents a selective force acting on modifiers of the corresponding mutation rate. This raises interesting perspectives on the joint evolutionary dynamics of non-coding genome size and structural mutation rates, which would certainly deserve further theoretical exploration.

To conclude, our results show that indirect selection against mutational hazards, *i*.*e*. robustness selection, and differences in the neutrality of mutations increasing or decreasing genome size are sufficient to explain the existence of an equilibrium in non-coding genome size. Structural mutations are sufficient to fulfill both conditions. The non-coding genome is constantly under indirect selection due to its mutagenic nature and the structural mutations it can initiate following double-strand breaks. As a consequence, a major determinant of the non-coding genome fraction is the product *N*_*e*_*× µ*, which affects both the efficacy of selection and the robustness cost of each additional base pair. More research should be conducted into that area to reach quantitative results and to understand more precisely how each determinant of non-coding genome size (*N*, *µ*, mutation bias, types of mutations, number and length of genes) affects the equilibrium non-coding size of a species, and whether the species are at that equilibrium or tending towards it. Finally, the interaction of indirect selection on the non-coding genome and direct selection on the coding genome should be studied further by relaxing the hypotheses of a fixed coding architecture and a binary fitness.

## Materials and Methods

### Logic behind the model

We consider a model with a simple genome architecture: organisms own a single circular chromosome. Mutations happen at random on the chromosome, each position for a mutation being drawn from a uniform distribution along the genome. While very simplistic, this approach carries an essential property of structural mutations: their size grows with total genome size [16]. This is easily demonstrated by biological data, as some observed inversions of several Mb [44] are bigger than the genomes of other organisms. The fact that bigger genomes are more susceptible to mutating and that structural mutations are increasingly dangerous implies a selection for shorter genomes on the lineage level, which we quantify mathematically.

The logic behind the model is the following (detailed computations are provided in the Supplementary Materials): the computation of each *ν*_*i*_ is made assuming a uniform draw of all positions needed for the mutation (two for deletions and three for duplications). Each mutation that has any phenotypic effect (deleting part of a gene, duplicating inside a gene, or duplicating a promoter) is lethal. To compute the effective fitness, we consider the probability for each base pair to initiate each type of mutation, and for the initiated mutation to be neutral. This corresponds to a binomial law where success is defined as either no mutation occurs or a neutral mutation occurs, the number of draws is the genome length multiplied by the number of mutation types. A non-lethal reproduction means there are only successes. The fixation probability is taken from Sella and Hirsh 2005 [39]. The average contribution to genome size change of each mutation type is computed as an average of the mutation size weighted by the probability of the mutation to be of this size and by the probability of a neutral mutation of this size to be fixed.

### Numerical resolution

All the code for the numerical resolution of the mathematical model is available online in GitLab: https://gitlab.inria.fr/jluisell/structural-mutations-set-an-equilibrium-non-coding-fraction. Notably, the code is written in `Python` using `Decimal` to increase float precision. This is necessary due to the very large values taken by our parameters (especially genome size and population size) and the several exponents in the computation.

To predict an equilibrium non-coding genome size, we fix the number of genes *g*, the coding genome size *z*_c_, the effective population size *N*_*e*_, the per-base per mutation type mutation rate *µ*, and the mutational bias *κ*. We then compute the resulting bias towards adding or losing bases for several possible non-coding sizes. Since the bias function is monotonous with respect to the non-coding genome size, we find the actual equilibrium with a bisect method.

### Biological data

To compare our model to actual biological data, we gathered mutation rates and effective population sizes from [26]. For each available species, we downloaded the annotated genome from NCBI and isolated the biggest chromosome. We counted any base pais annotated as part of a protein-coding gene as a “coding” base pair (*z*_c_), all the other base pairs as “non-coding” (*z*_nc_), and we counted the number of continuous non-coding segments (*g*). While more precise annotations could be used (*e*.*g*. regulatory RNAs should be counted as coding in our model, since they are functional), this approach reduced the annotation bias between well-studied model species and less annotated pioneer species.

## Acknowledgemnts

The authors thank Laurent Duret, Julien Joseph, Ivan Junier, and Flora Gaudillère for fruitful discussions on the model and the biological data. Funding: Agence Nationale de la Recherche (ANR-20-CE02-0008-01 NeGA).

