## Supplementary Materials for "Structural mutations set an equilibrium non-coding genome fraction"

#### Supplementary Information

#### Supplementary Note 1: Probability for a mutation to be neutral

For each type of mutation, we note $\nu_{\mathrm{mutation}}$ its probability to be perfectly neutral regarding the viability of the individual. We note $p_1$ the first position uniformly drawn on the genome, and $p_2$ and $p_3$ the second and third when needed. As such, each base has a probability $\frac{1}{L}$ to be drawn. There are $z_{\mathrm{nc}}$ non-coding bases, distributed along $g$ non-coding segments. The computations are provided for 6 types of mutations: deletions and duplications as well as point mutations, small insertions, small deletions, and inversions.

##### A. Probability for deletions to be neutral

A deletion is neutral if, and only if, the bases deleted are within one of the $g$ non-coding segments. This means that if the first deleted base is at a position $i$, $i$ must be in the non coding part of the genome, and the second must be at a position $j$ in the same non-coding region.

$$\begin{aligned}\nu_{\mathrm{del}}(g, z_{\mathrm{c}}, z_{\mathrm{nc}}) &= g \sum_{i=1}^{z_{\mathrm{nc}}/g} \left( \frac{1}{L} \sum_{j=i}^{z_{\mathrm{nc}}/g} \frac{1}{L} \right) \\ &= \frac{g}{2L^2} \sum_{i=1}^{z_{\mathrm{nc}}/g} \left( \frac{z_{\mathrm{nc}}}{g} - i + 1 \right) \\ &= \frac{z_{\mathrm{nc}} \left( \frac{z_{\mathrm{nc}}}{g} + 1 \right)}{2L^2}\end{aligned}$$

##### B. Probability for duplications to be neutral

A duplication is neutral if, and only if, it duplicates a sequence without a promoter basis and copy it at any position in the non-coding regions. The sum over $i$ starts at position 2 to avoid the first base (promoter), and then all duplications are valid as long as they do not encompass the next promoter. This probability is then multiplied by the probability for the insertion point to be in a non-coding region. Note that there are $z_{\mathrm{nc}}/g + 1$ insertion points in a non-coding sequence of size $z_{\mathrm{nc}}/g$ as we can insert just before and just after the sequence.

$$\begin{aligned}\nu_{\mathrm{dupl}}(g, z_{\mathrm{c}}, z_{\mathrm{nc}}) &= g \sum_{i=2}^{L/g} \left( \frac{1}{L} \sum_{j=i}^{L/g} \frac{1}{L} \right) \left( g \sum_{k=0}^{z_{\mathrm{nc}}/g} \frac{1}{L} \right) \\ &= \frac{g^2}{L^3} \sum_{i=2}^{L/g} \sum_{j=i}^{L/g} \left( \frac{z_{\mathrm{nc}}}{g} + 1 \right) \\ &= \frac{g(z_{\mathrm{nc}} + g)}{L^3} \sum_{i=2}^{L/g} \left( \frac{L}{g} - i + 1 \right) \\ &= \frac{g(z_{\mathrm{nc}} + g)}{L^3} \sum_{i=1}^{L/g-1} \left( \frac{L}{g} - i \right) \\ &= \frac{g(z_{\mathrm{nc}} + g) \left( \frac{L}{g} - 1 \right) \left( \frac{L}{g} \right)}{2L^3} \\ &= \frac{(z_{\mathrm{nc}} + g) \left( \frac{L}{g} - 1 \right)}{2L^2}\end{aligned}$$

##### C. Probability for point mutations to be neutral

Point mutations are neutral when they affect a non-coding base, and deleterious when they affect a coding base. The probability to affect a non-coding base is $\frac{z_{\mathrm{nc}}}{L}$:

$$\nu_{\mathrm{pm}}(g, z_{\mathrm{c}}, z_{\mathrm{nc}}) = g \sum_{i=1}^{z_{\mathrm{nc}}/g} \frac{1}{L} = \frac{z_{\mathrm{nc}}}{L}$$

##### D. Probability for small insertions to be neutral

Regardless of their size, small insertions are neutral when outside a coding segment, and deleterious when within a coding segment. Note however that there are $z_{\mathrm{nc}}/g+1$ insertion points in a non-coding sequence of size $z_{\mathrm{nc}}/g$ as we can insert just before and just after the sequence.

$$\nu_{\mathrm{indel}^+}(g, z_{\mathrm{c}}, z_{\mathrm{nc}}) = g \sum_{i=0}^{z_{\mathrm{nc}}/g} \frac{1}{L} = \frac{(z_{\mathrm{nc}}+g)}{L}$$

##### E. Probability for small deletions to be neutral

The maximum size $l_m$ of indels events is a parameter of the model. Here, we assume that $z_{\mathrm{nc}}/g \geq l_m$. The rationale here is to calculate the probability of a neutral deletion by separating all non-coding sequences into the $z_{\mathrm{nc}}/g-(l_m-1)$ first bases that can witness deletions of size up to $l_m$, and the $l_m-1$ other bases for which only a subset of the possible deletions are neutral. Since the length of the deletion is uniformly chosen between $1$ and $l_m$, if the deletion starts from a basis $i$ close to the end $z_{\mathrm{nc}}/g$ of the non coding zone, the probability that it is neutral is $\sum_{k=i}^{z_{\mathrm{nc}}/g} \frac{1}{l_m}$ when $z_{\mathrm{nc}}/g-i<l_m$

$$\begin{aligned}
\nu_{\mathrm{indel}^-}(g, z_{\mathrm{c}}, z_{\mathrm{nc}}) &= g\left(\sum_{i=1}^{z_{\mathrm{nc}}/g-(l_m-1)} \frac{1}{L} + \sum_{i=z_{\mathrm{nc}}/g-(l_m-2)}^{z_{\mathrm{nc}}/g} \frac{1}{L} \sum_{k=i}^{z_{\mathrm{nc}}/g} \frac{1}{l_m}\right) \\
&= \frac{g}{L}\left(\frac{z_{\mathrm{nc}}}{g} - (l_m-1) + \frac{1}{l_m} \sum_{z_{\mathrm{nc}}/g-(l_m-2)}^{z_{\mathrm{nc}}/g} \frac{z_{\mathrm{nc}}}{g} - i + 1\right) \\
&= \frac{1}{L}\left(z_{\mathrm{nc}} - g(l_m-1) + \frac{g}{l_m} \sum_{i=1}^{l_m-1} i\right) \\
&= \frac{1}{L}\left(z_{\mathrm{nc}} - g\frac{l_m-1}{2}\right)
\end{aligned}$$

##### F. Probability for inversions to be neutral

An inversion is neutral if the two breakpoints are outside coding regions. Note that the second breakpoint must be different from the first for an inversion to occur. The probability of the inversion to be neutral is thus the product of the two probabilities.

$$\begin{aligned}
\nu_{\mathrm{inv}}(g, z_{\mathrm{c}}, z_{\mathrm{nc}}) &= \frac{(z_{\mathrm{nc}}+g)}{L} \times \frac{(z_{\mathrm{nc}}+g)-1}{L-1} \\
&= \frac{(z_{\mathrm{nc}}+g)(z_{\mathrm{nc}}+g-1)}{L(L-1)}
\end{aligned}$$

#### Supplementary Note 2: Expected contribution to genome size change along evolution

For each mutation changing the genome size, we can compute its expected contribution per generation to the average genome size change for a species in terms of base pairs.

##### A. Contribution of deletions to genome size change

This corresponds to the average size of a deletion weighted by the probability of neutrality times the probability of fixation.

$$\begin{aligned}
\delta_{\mathrm{del}}(\mu, N, g, z_{\mathrm{c}}, z_{\mathrm{nc}}) &= g \sum_{i=1}^{z_{\mathrm{nc}}/g} \left( \frac{1}{L} \sum_{j=i}^{z_{\mathrm{nc}}/g} \frac{1}{L} (j-i+1) \mathbb{P}_{fix}(-(j-i+1)) \right) \\
&= \frac{g}{L^2} \sum_{i=1}^{z_{\mathrm{nc}}/g} \sum_{j=i}^{z_{\mathrm{nc}}/g} (j-i+1) \mathbb{P}_{fix}(-(j-i+1)) \\
&= \frac{g}{L^2} \sum_{i=1}^{z_{\mathrm{nc}}/g} \sum_{k=1}^{z_{\mathrm{nc}}/g-i+1} k \mathbb{P}_{fix}(-k) \\
&= \frac{g}{L^2} \sum_{k=1}^{z_{\mathrm{nc}}/g} \sum_{i=1}^{z_{\mathrm{nc}}/g-k+1} k \mathbb{P}_{fix}(-k) \\
&= \frac{g}{L^2} \sum_{k=1}^{z_{\mathrm{nc}}/g} \left( \frac{z_{\mathrm{nc}}}{g} - k + 1 \right) k \mathbb{P}_{fix}(-k) \\
&= \frac{1}{L^2} \sum_{k=1}^{z_{\mathrm{nc}}/g} (z_{\mathrm{nc}} - g(k+1)) k \mathbb{P}_{fix}(-k)
\end{aligned}$$

##### B. Contribution of duplications to genome size change

Similarly, this corresponds to the average size of a duplication weighted by the probability of neutrality times the probability of fixation.

$$\begin{aligned}
\delta_{\mathrm{dupl}}(\mu, N, g, z_{\mathrm{c}}, z_{\mathrm{nc}}) &= g \sum_{i=2}^{L/g} \left( \frac{1}{L} \sum_{j=i}^{L/g} \frac{1}{L} \big( g \sum_{k=0}^{z_{\mathrm{nc}}/g} \frac{1}{L} (j-i+1) \mathbb{P}_{fix}(j-i+1) \big) \right) \\
&= \frac{g(z_{\mathrm{nc}}+g)}{L^3} \sum_{i=2}^{L/g} \sum_{j=i}^{L/g} (j-i+1) \mathbb{P}_{fix}(j-i+1) \\
&= \frac{g(z_{\mathrm{nc}}+g)}{L^3} \sum_{i=2}^{L/g} \sum_{j=1}^{L/g-i+1} j \mathbb{P}_{fix}(j) \\
&= \frac{g(z_{\mathrm{nc}}+g)}{L^3} \sum_{j=1}^{L/g-1} \sum_{i=2}^{L/g-j+1} j \mathbb{P}_{fix}(j) \\
&= \frac{g(z_{\mathrm{nc}}+g)}{L^3} \sum_{j=1}^{L/g-1} \left( \frac{L}{g} - j \right) j \mathbb{P}_{fix}(j)
\end{aligned}$$

#### C. Contribution of small insertions (InDel$^+$) to genome size change

This corresponds to the average size of a small insertion weighted by its probability of neutrality times its probability of fixation.

$$\begin{aligned}\delta_{\text{indel}^+}(\mu, N, g, z_\text{c}, z_\text{nc}) &= g \sum_{i=0}^{z_\text{nc}/g} \frac{1}{L} \sum_{k=1}^{l_m} \frac{k\mathbb{P}_{fix}(k)}{l_m} \\ &= \frac{(z_\text{nc}+g)}{L\, l_m} \sum_{k=1}^{l_m} k\mathbb{P}_{fix}(k)\end{aligned}$$

#### D. Contribution of small deletions (InDel$^-$) to genome size change

This corresponds to the average size of a small deletion weighted by its probability of neutrality times its probability of fixation.

$$\begin{aligned}\delta_{\text{indel}^-}(\mu, N, g, z_\text{c}, z_\text{nc}) &= g \left( \sum_{i=1}^{z_\text{nc}/g-(l_m-1)} \left( \frac{1}{L} \sum_{k=1}^{l_m} \frac{k\mathbb{P}_{fix}(-k)}{l_m} \right) + \sum_{i=z_\text{nc}/g-(l_m-2)}^{z_\text{nc}/g} \frac{1}{L} \sum_{k=i}^{z_\text{nc}/g} \frac{(k-i+1)\mathbb{P}_{fix}(-(k-i+1))}{l_m} \right) \\ &= \frac{g}{L\, l_m} \left( \left( \frac{z_\text{nc}}{g} - (l_m-1) \right) \sum_{k=1}^{l_m} k\mathbb{P}_{fix}(-k) + \sum_{i=z_\text{nc}/g-(l_m-2)}^{z_\text{nc}/g} \sum_{j=1}^{z_\text{nc}/g-i+1} j\mathbb{P}_{fix}(-j) \right) \\ &= \frac{1}{L\, l_m} \left( (z_\text{nc} - g(l_m-1)) \sum_{k=1}^{l_m} k\mathbb{P}_{fix}(-k) + \sum_{s=1}^{l_m-1} \sum_{j=1}^{s} j\mathbb{P}_{fix}(-j) \right)\end{aligned}$$

#### Supplementary Note 3: Joined impact of $N$ and $\mu$ on non-coding genome fraction at equilibrium

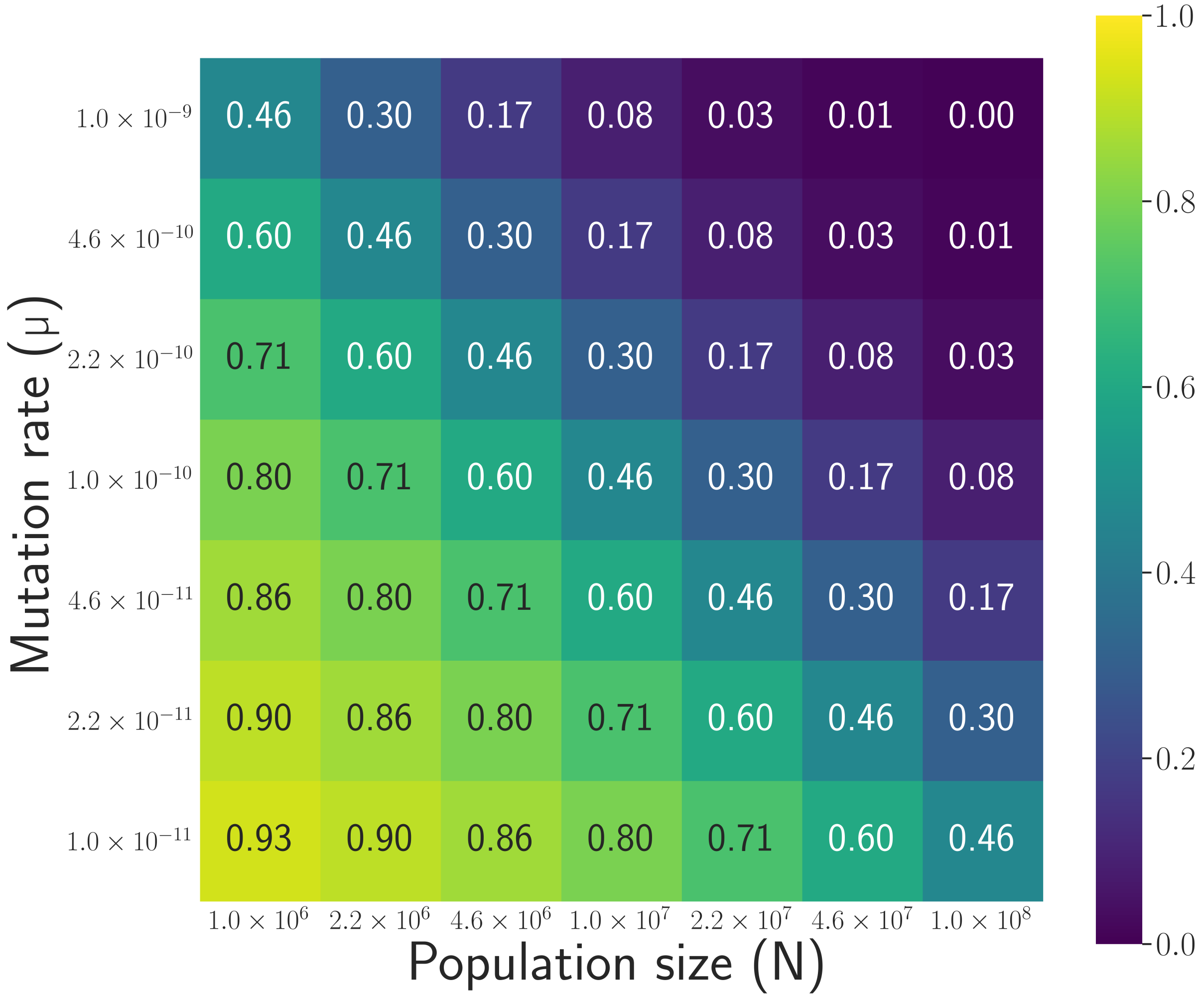


**Figure S1.** Predicted non-coding fraction at equilibrium for different values of $N$ and $\mu$. The genome architecture is fixed at $z_c = 1,000,000$ and $g = 2,000$, and we have $\lambda_{\text{dupl}} = \lambda_{\text{del}} = 1$.

A change in $N$ or in $\mu$ by the same factor results in the same non-coding fraction.

#### Supplementary Note 4: Simplified model with only indels (no structural mutations)

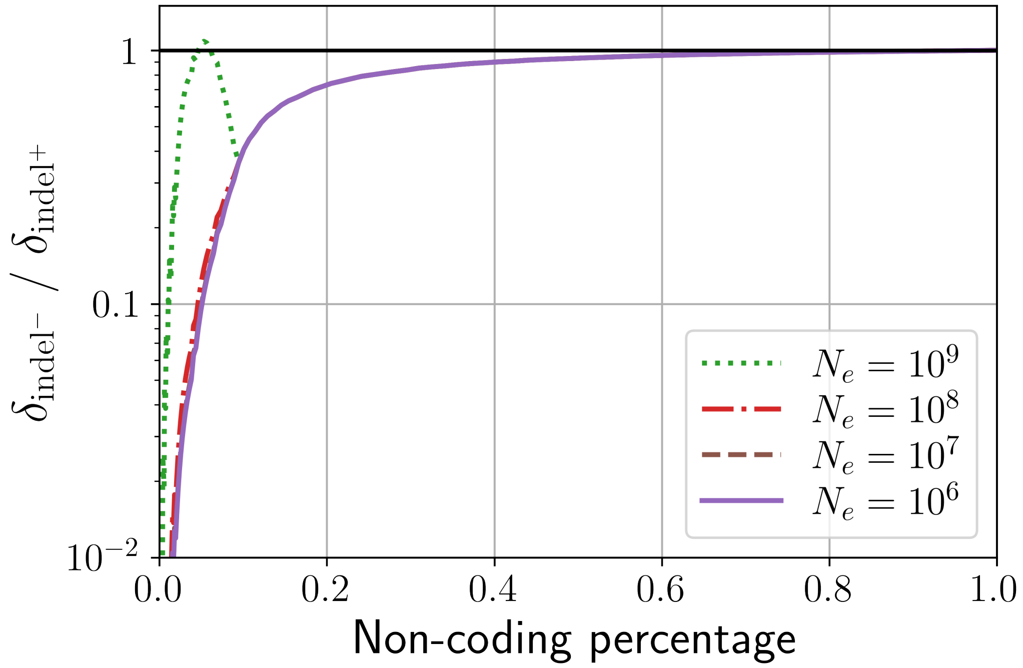


**Figure S2.** Measured bias for different non-coding proportions. Genome architecture is fixed at $z_c = 1{,}000{,}000$ and $g = 2{,}000$, the mutation rate is fixed at $\mu = 1 \times 10^{-10}$ and $\lambda_{\text{indel}^-} = \lambda_{\text{indel}^+} = 1$. The maximum size of indels ($l_m$) is $50$. $z_{\text{nc}}$ varies in a logspace from $10^3$ to $10^9$, and four different values of $N$ are depicted. The black horizontal line shows the potential equilibrium at $B = 1$, but is only crossed for $N_e = 10^9$.

Except for the highest value of $N_e$, the bias converges towards $1$ as the non-coding proportion increases but is always below $1$: when only indels are modeled, genome size would grow indefinitely in most cases in our model. Yet, when non-coding segments of the genome are small compared to the size of indels ($z_{nc}/g \ll l_m$), small deletions create a selection for robustness similar to the one of structural mutations: indels are more numerous as genome size increases but only marginally less deleterious and so genome growth could counter-selected. As a result, if the selection for robustness is very strong (very large $N_e$), it can counterbalance the difference in neutrality between small insertions and small deletions and thus there can be an equilibrium genome size, at a size where the intergenic segments are lower than the maximal size of indels.

#### Supplementary Note 5: Equations with the full set of mutations

We note $M$ the set of six mutations: duplications, deletions, inversions, small insertions, small deletions, and point mutations.

##### A. Effective fitness

$$f_e(\mu, g, z_c, z_{\text{nc}}) = \prod_{i \in M} \left(1 - \mu + \mu\, \nu_i(g, z_c, z_{\text{nc}})\right)^L \tag{S1}$$

##### B. Overall bias

$$\begin{aligned} B(\mu, N, g, z_c, z_{\text{nc}}) &= \frac{\mu\, L\, N\, \delta_{\text{del}}(\mu, N, g, z_c, z_{\text{nc}}) + \mu\, L\, N\, \delta_{\text{indel}^-}(\mu, N, g, z_c, z_{\text{nc}})}{\mu\, L\, N\, \delta_{\text{dupl}}(\mu, N, g, z_c, z_{\text{nc}}) + \mu\, L\, N\, \delta_{\text{indel}^+}(\mu, N, g, z_c, z_{\text{nc}})} \\ &= \frac{\delta_{\text{del}}(\mu, N, g, z_c, z_{\text{nc}}) + \delta_{\text{indel}^-}(\mu, N, g, z_c, z_{\text{nc}})}{\delta_{\text{dupl}}(\mu, N, g, z_c, z_{\text{nc}}) + \delta_{\text{indel}^+}(\mu, N, g, z_c, z_{\text{nc}})} \end{aligned} \tag{S2}$$

### Supplementary Note 6: Average size of spontaneous mutations

We want to compute the spontaneous contribution of the different mutations to genome size changes. As non-neutral mutations are lethal, their size is counted as 0: they cannot change the genome size. Four types of mutations can change the genome size: deletions and duplications, as well as small deletions and small insertions.

##### A. Genome size change due to a neutral deletion

$$\begin{aligned}
\eta_{del} &= g \sum_{i=1}^{z_{\mathrm{nc}}/g} \left( \frac{1}{L} \sum_{j=i}^{z_{\mathrm{nc}}/g} \frac{1}{L}(j-i+1) \right) \\
&= \frac{g}{L^2} \sum_{i=1}^{z_{\mathrm{nc}}/g} \sum_{j=i}^{z_{\mathrm{nc}}/g} (j-i+1) \\
&= \frac{g}{L^2} \sum_{i=1}^{z_{\mathrm{nc}}/g} \sum_{j=1}^{z_{\mathrm{nc}}/g-i+1} j \\
&= \frac{g}{2L^2} \sum_{i=1}^{z_{\mathrm{nc}}/g} (z_{\mathrm{nc}}/g-i+1)(z_{\mathrm{nc}}/g-i+2) \\
&= \frac{z_{\mathrm{nc}}(\frac{z_{\mathrm{nc}}}{g}+1)(\frac{z_{\mathrm{nc}}}{g}+2)}{6L^2}
\end{aligned}$$

##### B. Genome size change due to a neutral duplication

$$\begin{aligned}
\eta_{\mathrm{dupl}}(g, z_{\mathrm{c}}, z_{\mathrm{nc}}) &= g \sum_{i=2}^{L/g} \left( \frac{1}{L} \sum_{j=i}^{L/g} \frac{1}{L} \left( g \sum_{k=0}^{z_{\mathrm{nc}}/g} \frac{1}{L}(j-i+1) \right) \right) \\
&= \frac{g(z_{\mathrm{nc}}+g)}{L^3} \sum_{i=2}^{L/g} \sum_{j=i}^{L/g} (j-i+1) \\
&= \frac{g(z_{\mathrm{nc}}+g)}{L^3} \sum_{i=2}^{L/g} \sum_{j=1}^{L/g-i+1} j \\
&= \frac{g(z_{\mathrm{nc}}+g)}{2L^3} \sum_{i=2}^{L/g} \left( \frac{L}{g} - i + 1 \right) \left( \frac{L}{g} - i + 2 \right) \\
&= \frac{g(z_{\mathrm{nc}}+g)}{2L^3} \sum_{i=1}^{L/g-i} (i)(i+1) \\
&= \frac{g(z_{\mathrm{nc}}+g) \left( \frac{L}{g} - 1 \right) \left( \frac{L}{g} \right) \left( \frac{L}{g} + 1 \right)}{6L^3} \\
&= \frac{(z_{\mathrm{nc}}+g) \left( \frac{L}{g} - 1 \right) \left( \frac{L}{g} + 1 \right)}{6L^2}
\end{aligned}$$

##### C. Genome size change due to a neutral Indel$^-$

A small deletion has size $k \leq l_m$ with probability $\frac{1}{l_m}$ so the mean size is :

$$
\begin{aligned}
\eta_{\text{indel}^-}(g, z_c, z_{nc}) &= g\left(\sum_{i=1}^{z_{nc}/g-(l_m-1)}\left(\frac{1}{L}\sum_{k=1}^{l_m}\frac{k}{l_m}\right) + \sum_{i=z_{nc}/g-(l_m-2)}^{z_{nc}/g}\frac{1}{L}\sum_{k=i}^{z_{nc}/g}\frac{(k-i+1)}{l_m}\right)\\
&= \frac{g}{L\,l_m}\left(\sum_{i=1}^{z_{nc}/g-(l_m-1)}\frac{l_m(l_m+1)}{2} + \sum_{i=z_{nc}/g-(l_m-2)}^{z_{nc}/g}\sum_{j=1}^{z_{nc}/g-i+1} j\right)\\
&= \frac{g}{L\,l_m}\left(\frac{l_m(l_m+1)}{2}\left(\frac{z_{nc}}{g}-(l_m-1)\right) + \sum_{i=z_{nc}/g-(l_m-2)}^{z_{nc}/g}\frac{\left(\frac{z_{nc}}{g}-i+1\right)\left(\frac{z_{nc}}{g}-i+2\right)}{2}\right)\\
&= \frac{1}{2L}\left((l_m+1)(z_{nc}-g(l_m-1)) + \sum_{j=1}^{l_m-1} j(j+1)\right)\\
&= \frac{1}{2L}\left((l_m+1)(z_{nc}-g(l_m-1)) + \frac{l_m(l_m-1)(2l_m-1)}{6} + \frac{l_m(l_m-1)}{2}\right)\\
&= \frac{1}{2L}\left(z_{nc}(l_m+1) - g({l_m}^2-1) + \frac{2{l_m}^2-3l_m+1}{6} + \frac{l_m-1}{2}\right)\\
&= \frac{1}{L}\left(z_{nc}\frac{l_m+1}{2} + g\frac{1-{l_m}^2}{3}\right)
\end{aligned}
$$

##### D. Genome size change due to a neutral Indel$^+$

$$
\begin{aligned}
\eta_{\text{indel}^+}(g, z_c, z_{nc}) &= g\sum_{i=0}^{z_{nc}/g}\frac{1}{L}\sum_{k=1}^{l_m}\frac{k}{l_m}\\
&= \frac{g}{L\,l_m}\left(\sum_{i=0}^{z_{nc}/g} 1\right)\left(\sum_{k=1}^{l_m} k\right)\\
&= \frac{(z_{nc}+g)(l_m+1)}{2L}
\end{aligned}
$$

After calculation, we have $\eta_{\text{indel}^+} > \eta_{\text{indel}^-}$, and since $z_c/g \geq 1$ unless all coding sections are only composed of promoter sequences, we also have $\eta_{\text{dupl}} > \eta_{\text{del}}$. Thus, there exists a neutral bias towards non coding genome size increase. If phenotypical adaptation was constant, genomes would tend to gain more new non-coding bases through duplications than what they lose through deletions. However, we do not observe an infinite growth of genome sizes, and that is due to a variation in the probability of fixation of our neutral mutations.
